# Establishment, verification and application of rapid detection of baculovirus infectious titer by flow cytometry

**DOI:** 10.1101/2021.05.26.445796

**Authors:** Wu Qingsheng, Li Yuanyuan

**Affiliations:** No.7 Research Laboratory, National Vaccine and Serum Institute, Beijing 101111, China

**Author notes:** Corresponding author: Li Yuanyuan.

**Keywords:** Flow cytometry, Baculovirus titer, Titer detection

## Abstract

Titer detection of baculovirus usually is time-consuming.It is important to establish a rapid detection method for baculovirus titer. In this report, Staining of cells with a fluorescently labeled anti-gp64 antibody allows for identification of infected insect cells. By inoculating cultures with a series of log dilutions of virus, and staining of the cultures 13-22 hours post inoculation, the ratio of infected to un-infected insect cells can be determined by flow cytometry. Statistical analysis of the percentage of infected cells in the virus dilution series enables accurate infectious titer determination. The culture time, cell growth state, the concentration of GP64-APC antibody and the concentration of inactivated FBS in diluent were optimized.The generality, repeatability and intermediate precision of the method were verified. The FCM method has the advantages of simplicity, accuracy, low cost and good repeatability.

## Introduction

Baculovirus expression vector system (Baculovirus expression vector system, BEVS) can express foreign genes at a high level. The biological characteristics of the expression product are similar to those of natural products. It has good application prospects in vaccine and drug development. Seven products including cancer vaccines and influenza vaccines have been approved, including vaccines and therapeutic products^[1]^. It is worth mentioning that on November 16, 2020, the new coronavirus recombinant protein vaccine developed by West China Hospital of Sichuan University using insect cells has entered the phase II clinical stage^[2]^.

The determination of baculovirus titer is very important for the control and optimization of recombinant protein expression process parameters. Baculovirus determination methods mainly include plaque method, endpoint dilution method (TCID50), live cell size determination method, fluorescent quantitative PCR method, flow cytometry detection method, immunostaining method, colorimetric indicator, and β-half determination. Lactosidase activity, the use of microfluidic bioanalyzer, etc^[3]^. The plaque method is a classic method for titer determination, but the detection cycle is long (6-10 days), and the operator’s technical and experience requirements are high; the end-point dilution method also requires at least 3 days. Currently, immunostaining and qPCR are widely used. The qPCR method has the advantage of a short detection cycle, but it requires a nucleic acid extraction step, which is to detect the number of gene copies, rather than the infectious virus titer value, which is not accurate enough. The immunostaining method also has a series of shortcomings: there are many operating steps and a long detection cycle; manual counting is required, which is more labor-intensive; insect cells are semi-adherent cells, which are not firmly attached during detection, and they may fall off in the multi-well plate. In the process of counting spots, the subjectivity is strong and requires certain experience or standards to distinguish.

The FCM method is a technique commonly used to identify and isolate specific cells from mixed samples^[4]^. It can be used to detect viruses and cells infected by viruses^[5]^. Gp64 protein locates in the cytoplasm in the early stage of virus infection, then migrates to the plasma membrane, and can be expressed on the surface of infected insect cells within 6 hours of infection^[6∼7]^. Refer to the relevant literature, combined with the actual application, stain the cells with fluorescently labeled anti-gp64-APC antibody, and establish an FCM method for rapid detection of baculovirus titer^[8]^, and may affect the titer detection The value of incubation time, cell state, antibody concentration and other factors have been optimized and verified.

## 1 Materials and methods

### 1.1 Viruses and cells

Recombinant H5N1-HA, NA, M1, M2, NP Baculovirus, Recombinant H1N1-HA Baculovirus, Recombinant HEV Baculovirus, Recombinant 2019-nCOV-N Baculovirus, 2019-nCOV-S Baculovirus, 2019 -nCOV-S1, RBD, RBD dimer baculovirus and empty baculovirus were prepared by the No.7 Research Laboratory, National Vaccine and Serum Institute; Sf9 insect cells were purchased from ATCC; ExpiSf9 insect cells were purchased from ThermoFisher Scientific;

### 1.2 Reagents and instruments

SFX-Insect medium was purchased from GE,; Baculovirus Envelope gp64 Monoclonal Antibody (AcV1), APC, and eBioscience™ were purchased from ThermoFisher Scientific; TC Plate 24 Well, Suspension, F were purchased from Sarstedt; Gibco fetal bovine serum was purchased from Thermo Scientific ; PBS and resuspension (including 2% FBS), self-prepared. FACSCalibur™ flow cytometer and Falcon flow cytometry tube were purchased from BD; 3-18K refrigerated centrifuge was purchased from Sigma; QT-2 vortex mixer was purchased from Shanghai Qite Analytical Instrument Co., Ltd.; ZWY -211C constant temperature culture oscillator was purchased from Shanghai Zhicheng Analytical Instrument Manufacturing Co., Ltd.; Countstat^®^ BioTech automatic cell counter was purchased from Shanghai Ruiyu Biotechnology Co., Ltd.;

### 1.3 Method establishment

Add 1ml logarithmically diluted HEV virus solution and 800μl to a 24-well suspension culture plate with a density of 1.25×10[6]cells logarithmic phase insect cells, set up multiple wells and negative control wells. 27°C shaker 225rpm, 27°C suspension culture for 14-16 hours, aspirate the liquid from each well to the flow cytometry tube and centrifuge to pellet the cells (300×g, 5 minutes), discard the supernatant, and add 100μl at a concentration of 0.15μg/ml Stain the cells with the gp64-APC antibody, vortex and mix for 3 to 5 seconds, let stand at room temperature for 30 minutes, centrifuge to discard the supernatant, slowly add 1ml PBS along the tube wall to wash, add 1ml resuspension solution to resuspend the cells. Flow cytometry detection, select the detection value of positive cells less than 10% to calculate the virus titer.

Calculation formula (Infectious virus particle, ivp: infectious virus particle):

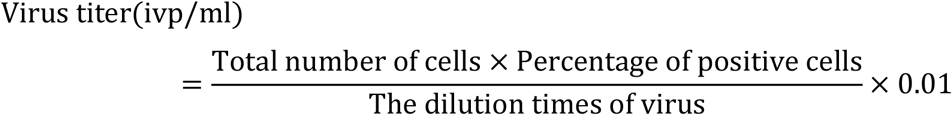

### 1.4 Method optimization

#### 1.4.1 incubation time

After 6-48 hours of baculovirus inoculation and culture, test to observe cell growth and titer change trends to determine the best culture time.

#### 1.4.2 Concentration of gp64 antibody

Add 100 μl of gp64-APC antibody with concentrations of 0.05, 0.1, 0.15, 0.2, 0.4 μg/ml for detection. Observe the trend of titer changes and select the best antibody concentration.

#### 1.4.3 Concentration of inactivated FBS in dilute diluent

Use HEV virus with 0%, 0.2%, 0.5%, 0.8%, 1%, 2%, 5%, 8%, 10% dilutions to resuspend stained cells washed with PBS and observe different FBS contents Whether the resuspension of the solution affects the detection value of titer, and choose the best FBS concentration.

#### 1.4.4 Cell growth status

1.4.4.1 Use cells of different growth periods (early, middle, late, plateau, and decay periods of logarithmic growth) and different cell generations for titer detection

1.4.4.2 Use another recombinant baculovirus: 2019-nCOV-RBD dimer-working virus library to verify the influence of cell growth status on the detection value.

### 1.5 Method verification

#### 1.5.1 versatility

Use the FCM method to detect baculoviruses carrying different foreign genes (H5N1-HA, H5N1-NA, H5N1-M1, H5N1-M2, H5N1-NP, H1N1-HA, 2019-nCOV-N, 2019-nCOV-S) Perform testing to verify the versatility of the method.

#### 1.5.2 Repeatability

For the same virus sample, repeat the test 6 times and calculate the RSD value.

#### 1.5.3 Intermediate precision

The same sample was repeatedly tested by three different operators to calculate the RSD value.

### 1.6 Method application

The FCM method was used to detect the titer of 2019-nCOV-S1, 2019-nCOV-RBD, and 2019-nCOV-RBD dimer in the tertiary virus seed library. The results were compared with the detect value of the immune-fluorescence method.

## 2 results

### 2.1 Method establishment

Use the P44 generation, 1 day after the passage of Sf9 cells, to detect the HEV virus, the detection values were 5.91×10^8^ ivp/ml, 5.68×10^8^ ivp/ml, the average value was 5.79 ×10^8^ ivp/ml.

### 2.2 Method optimization

#### 2.2.1 Incubation time

The cell and virus culture time is less than 13 hours, and the detection value is seriously low; between 13∼22 hours, the detection value is not much different; between 26∼32 hours, the detection value is reduced; between 36∼40 hours, the detection value has a slight upward trend; At 42 hours, the detection signal decreased significantly and almost disappeared; between 44∼48 hours, the detection value increased significantly again (Figure 2).

**Figure 1.**
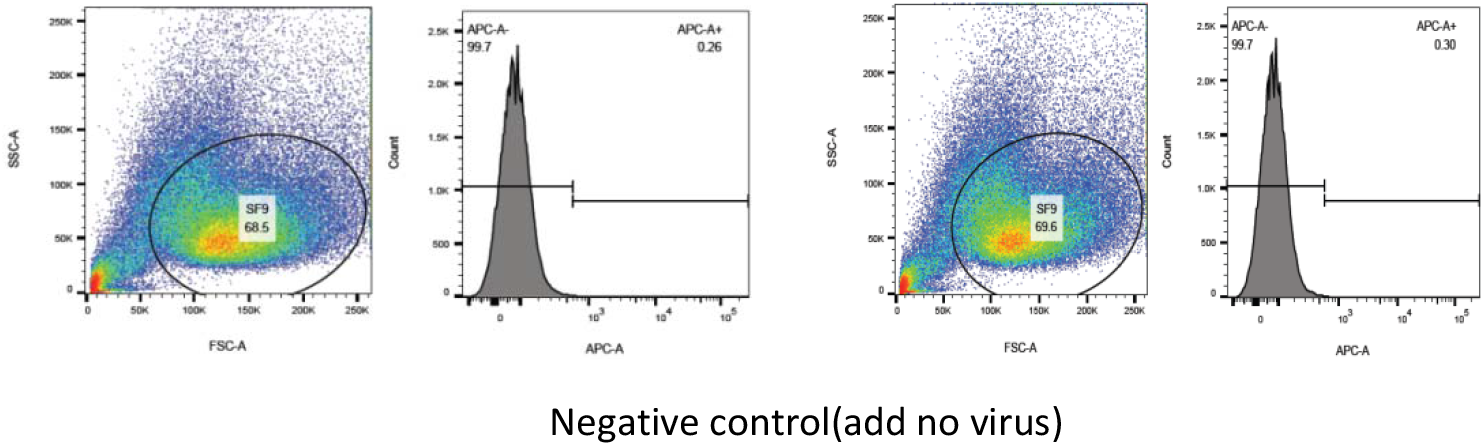
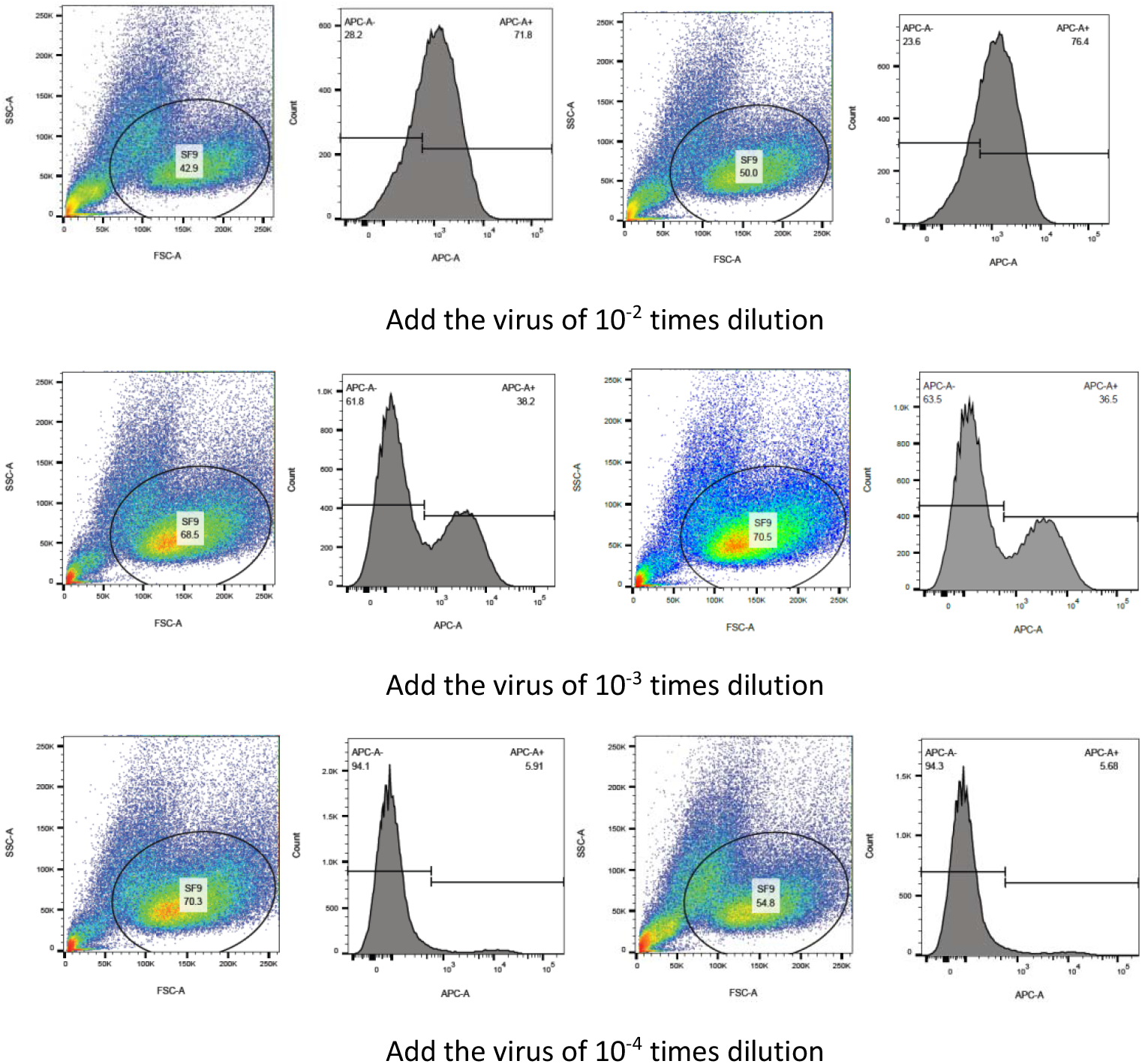
FCM scatter plots of Sf9 cells infected with different dilutions of virus

**Figure 2.**
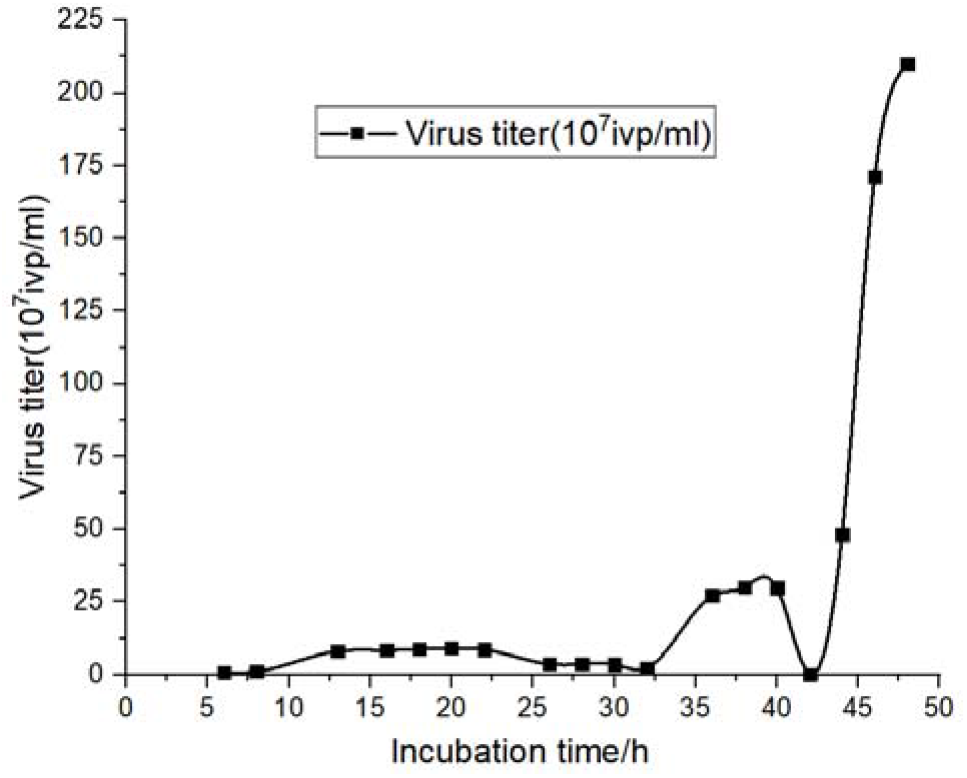
Changes of virus titer detection values at different incubation times

As the culture time increases, the cells that are not infected by the virus will continue to expand and the cell density will increase (Figure 3).

**Figure 3.**
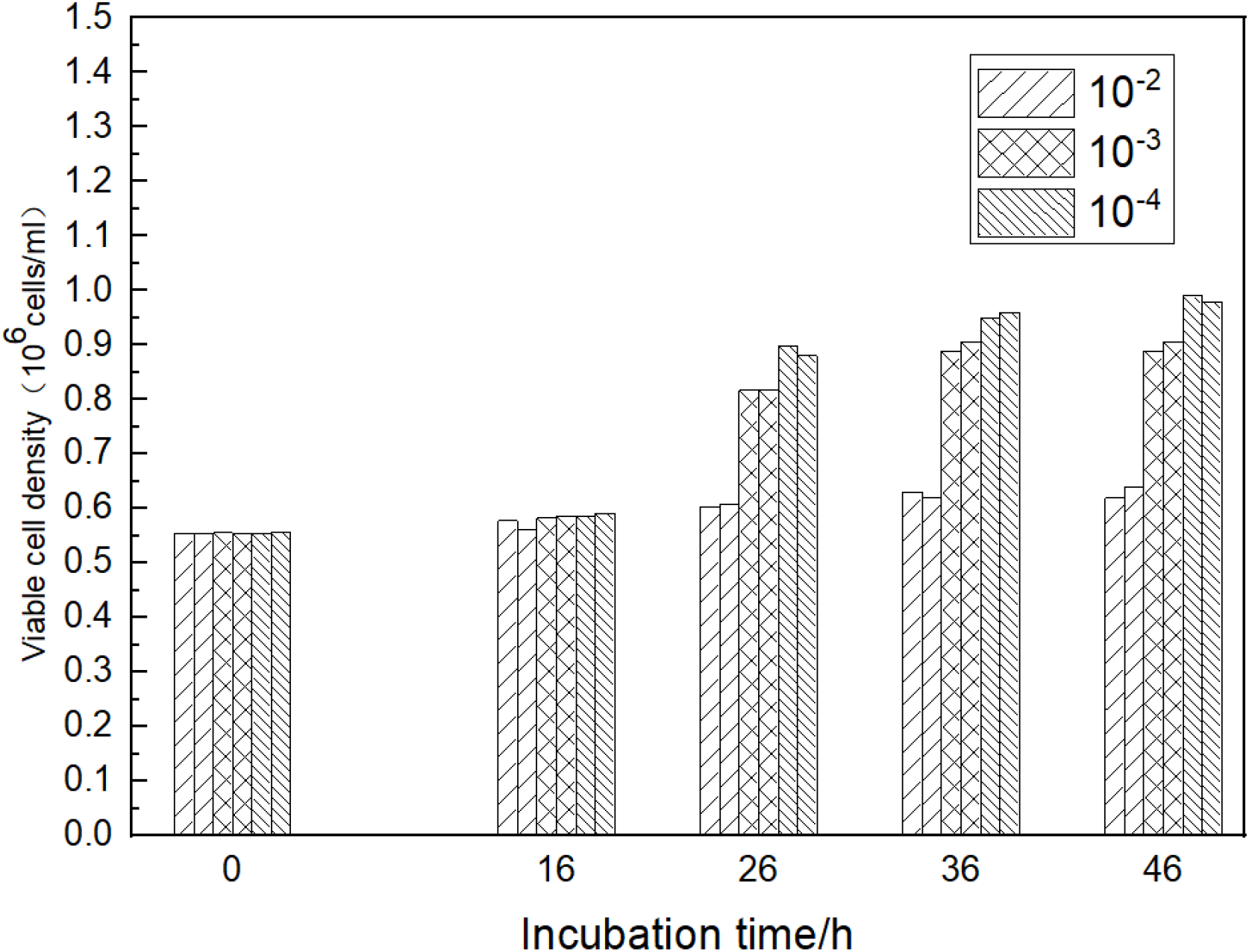
Changes of the total number of living cells infected with different culture time and different dilution ratio of virus

**Figure 4.**
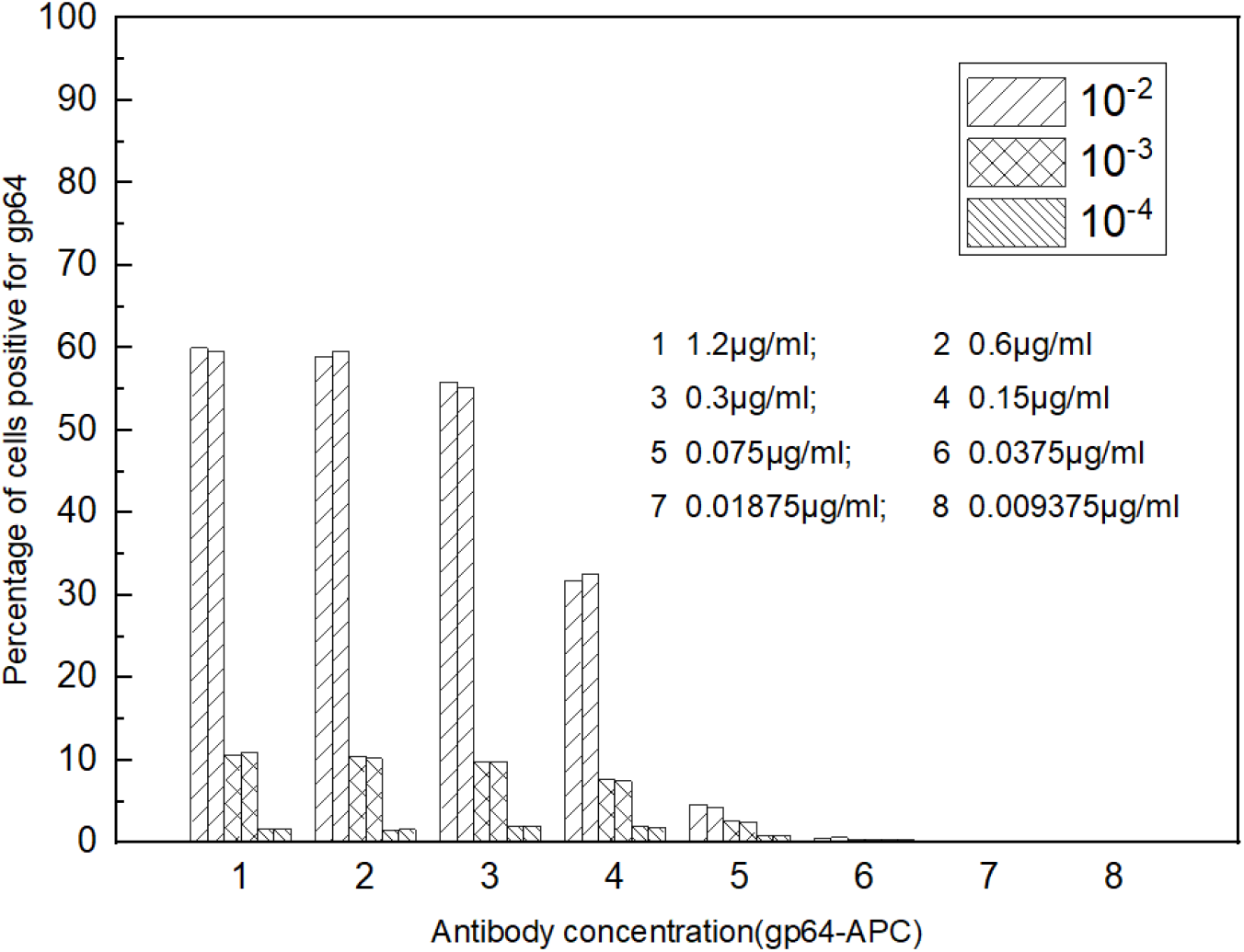
The effect of GP64-APC antibody concentration on virus titer detection value

**Figure 5.**
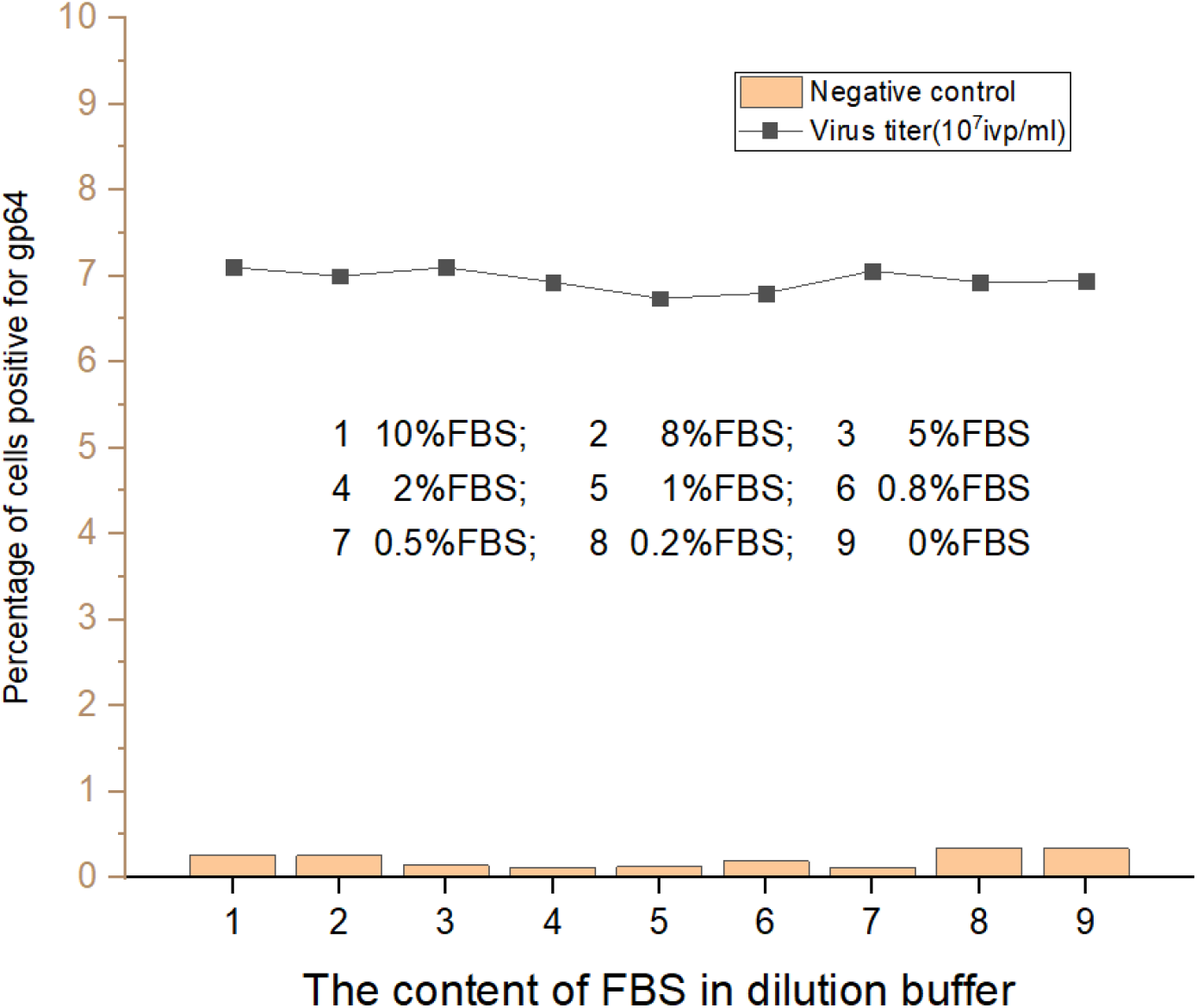
The effect of inactivated FBS content in diluents on detection value

#### 2.2.2 gp64-APC antibody concentration

The concentration of 100μl gp64-APC antibody for cell staining between 0.15∼1.2μg/ml has little effect on the detection value, and has no effect on the detection value of about 10% positive cell wells, so 0.15μg/ml is selected as the detection antibody concentration. If it is less than 0.15μg/ml, it will seriously affect the detection value; if it is less than 0.0375μg/ml, the proportion of positive cells in each dilution cannot be distinguished from the negative control.

#### 2.2.3 The content of inactivated FBS in the diluent

The content of inactivated FBS in the dilution has no significant effect on the detection value of HEV virus titer. The average detection value of 9 groups is 6.95, and the proportion of gp64 positive cells in the 2% FBS group is 6.93, which is close to the mean value; the negative control value between each group (Detection background) The difference is not obvious, the 2% concentration group is 0.11, which is relatively low. Therefore, the content of inactivated FBS in the diluent is 2%.

#### 2.2.4 Cell growth status

According to the detection value of Sf9 cell titer of different cell cycle stages (Table 1), it can be seen that the detection value of the cells in the early stage of logarithmic growth is higher and the stability is better; Use the cells in the **adaptive and** plateau phase, the detection value is lower; The cells in the recession cannot be used for titer detection due to the extremely poor cell state or low viability; The cell generation had no significant effect on the detection value of virus titer.High-passage cells (119 generations) are not suitable for titer detection;but, after 3 passages (122 generations), they can be used for titer detection again(Table 2).

**Table 1.** Comparison of titer values of Sf9 cells in different cell cycle stages

| Generation | Culture days | Viable cell density<br>(10<sup>6</sup>cells/ml) | viability (%) | diameter (μm) | Phase | Titer-1<br>(10<sup>7</sup>ivp/ml) | Titer-2<br>(10<sup>7</sup>ivp/ml) | Titer-Average<br>(10<sup>7</sup>ivp/ml) |
| --- | --- | --- | --- | --- | --- | --- | --- | --- |
| 40 | 2 | 2.97 | 97.51 | 16.98 | Early-logarithmic | 9 | 8.8 | 8.9 |
| 40 | 5 | 11.2 | 97.35 | 17.67 | plateau | 4.9 | 4.7 | 4.8 |
| 42 | 3 | 3.6 | 96.4 | 16.7 | Early-logarithmic | 8.3 | 8.1 | 8.2 |
| 44 | 1 | 1.4 | 95.35 | 16.53 | Early-logarithmic | 8.8 | 9 | 8.9 |
| 44 | 3 | 5.52 | 98.96 | 16.03 | Middle-logarithmic | 7.1 | 7.1 | 7.1 |
| 47 | 1 | 1.6 | 98.39 | 16.46 | adaptive | 7.7 | 7.5 | 7.6 |
| 47 | 2 | 3.5 | 98.71 | 16.28 | Early-logarithmic | 8.1 | 8.3 | 8.2 |
| 48 | 1 | 1.3 | 98.83 | 16.5 | adaptive | 7.5 | 7.1 | 7.3 |
| 54 | 7 | 3 | 29.8 | 15.55 | Recession | Not-suitable | Not-suitable | Not-suitable |
| 55 | 5 | 7.41 | 69.84 | 16.34 | recession | Not-suitable | Not-suitable | Not-suitable |
| 60 | 2 | 3.7 | 98.72 | 16.06 | Early-logarithmic | 8.4 | 8.2 | 8.3 |
| 119 | 4 | 8.92 | 90.58 | 17.57 | Later-logarithmic | Not-suitable | Not-suitable | Not-suitable |
| 122 | 1 | 2.78 | 92.5 | 17.02 | Early-logarithmic | 8.0 | 8.7 | 8.4 |

**Table 2.** Comparison of titer detection values of high passage cells (199 passage) and the cells which after three passage (122 passage)

| Sample | SF9/APC-A+ \| Freq. of Parent |  |
| --- | --- | --- |
|  | Sf9-P119-4 天 | Sf9-P122-1 天 |
| Negative control | 8.88% | 1.55% |
| 10<sup>-2</sup> times dilution | 25.4% | 40.4% |
| 10<sup>-3</sup> times dilution | 11.6 % | 8.01% |
| 10<sup>-3</sup> times dilution | 12.4 % | 8.77% |
| 10<sup>-4</sup> times dilution | 9.92 % | 2.23% |
| 10<sup>-4</sup> times dilution | 9.15 % | 2.85% |

**Table 3.**
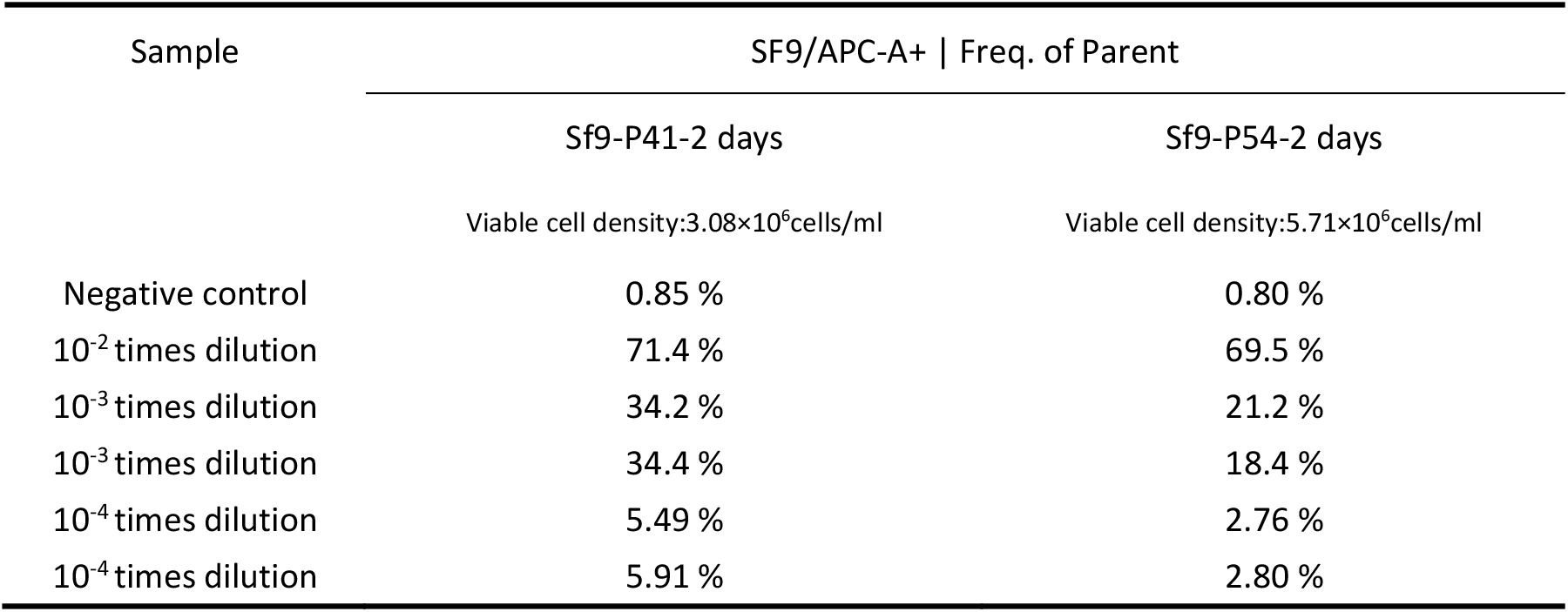
Another virus was used to verify the influence of cell status on titer detection values

**Table 4.** Universality verification

| Baculovirus | Titer-1<br>(10<sup>7</sup>ivp/ml) | Titer-2<br>(10<sup>7</sup>ivp/ml) | Titer-Average<br>(10<sup>7</sup>ivp/ml) |
| --- | --- | --- | --- |
| H5N1-HA | 49.40 | 53.60 | 51.50 |
| H5N1-NA | 4.50 | 4.38 | 4.44 |
| H5N1-M1 | 5.55 | 5.07 | 5.31 |
| H5N1-M2 | 4.13 | 4.18 | 4.16 |
| H5N1-NP | 2.86 | 2.88 | 2.87 |
| H1N1-HA | 16.8 | 17.60 | 17.20 |
| 2019-nCOV-N | 96.40 | 108.00 | 102.20 |
| 2019-nCOV-S | 58.10 | 58.70 | 58.40 |
| Empty baculovirus | 193.00 | 198.00 | 195.50 |

**Table 5.** Repeatability verification

| Times of repetition | Titer-1<br>($10^7$ivp/ml) | Titer-2<br>($10^7$ivp/ml) | Titer-Average<br>($10^7$ivp/ml) |
| --- | --- | --- | --- |
| 1 | 8.29 | 8.54 | 8.42 |
| 2 | 8.48 | 8.13 | 8.31 |
| 3 | 8.49 | 8.16 | 8.33 |
| 4 | 8.26 | 8.52 | 8.39 |
| 5 | 8.67 | 8.52 | 8.60 |
| 6 | 8.63 | 8.59 | 8.61 |
| Average |  |  | 8.44 |
| RSD (%) |  |  | 1.58 |

**Table 6.** The intermediate precision

| Times of repetition | The operator A-Titer<br>($10^7$ivp/ml) | The operator B-Titer<br>($10^7$ivp/ml) | The operator C-Titer<br>($10^7$ivp/ml) |
| --- | --- | --- | --- |
| 1 | 8.29 | 8.22 | 8.32 |
| 2 | 8.16 | 8.37 | 8.35 |
| 3 | 8.4 | 8.08 | 8.12 |
| Average | 8.28 | 8.22 | 8.26 |
| RSD (%) | 1.45 | 1.76 | 1.51 |
| RSD-Average (%) |  | 1.41% |  |

**Table 7.** FCM method and fluorescence immunoassay were used to detect the titer of 2019-nCOV virus

| Virus Bank | Titer-1<br>($10^7$ivp/ml) | Titer-2<br>($10^7$ivp/ml) | Titer-Average<br>($10^7$ivp/ml) | fluorescence<br>immunoassay($10^7$PFU/ml) |
| --- | --- | --- | --- | --- |
| S1-Primary | 6.99 | 7.18 | 7.09 | 7.51 |
| S1-Secondary | 28.70 | 26.10 | 27.40 | 38.60 |
| S1-Working | 45.40 | 46.50 | 45.95 | 32.85 |
| RBD-Primary | 16.60 | 13.30 | 14.95 | 18.42 |
| RBD-Secondary | 26.80 | 27.20 | 27.00 | 24.57 |
| RBD-Working | 36.60 | 33.30 | 34.95 | 28.62 |
| RBD dimer-Primary | 47.50 | 48.20 | 47.85 | 42.40 |
| RBD dimer -Secondary | 35.10 | 35.60 | 35.35 | 38.93 |
| RBD dimer -Working | 54.90 | 59.10 | 57.00 | 58.76 |

### 2.3 Method verification

#### 2.3.1 Versatility

The FCM method can detect baculoviruses carrying different foreign genes, and the method has a good versatility.

#### 2.3.2 Repeatability

The same virus sample was tested repeatedly for 6 times, and the calculated RSD=1.58%, with good repeatability.

#### 2.3.3 Intermediate precision

Three testers repeated the test for the same virus sample three times, with RSD-average=1.41%, with good intermediate precision.

### 2.4 Method application

Flow cytometry was used to detect the titers of the seed bank, RBD and RBD dimer of the 2019-NCoV-S1 virus. There was no significant difference between the results of flow cytometry and immune-fluorescence.

## 3 Discussion

The commercial kit (Expression Systems) of FCM method uses Gp64-PE antibody for cell staining, and the kit is restricted sales in some areas. In this article, Gp64-APC antibody is used for detection. The concentration of this antibody was 0.015μg/test, which was lower than that of GP64-PE antibody. The titer value is calculated according to the virus dilution well, which the positive cell proportion is about 10%, so there is no need to draw a standard curve, so it is more efficient.

Unlike the plaque method, the FCM method uses suspension culture, closely reflect actual culturing conditions utilized for expression cultures, and can obtain titer results within 24 hours. By using a centrifuge and a flow cytometer equipped to handle 96-well plates, the FCM method can easily achieve high-throughput detection. Since the FCM method detects the percentage of positive cells, it is not necessary to analyze all cells to get the correct detection value, during cell culture, centrifugation, cell staining, washing, etc., even if the cell count is inaccurate, or the total cell volume is lost due to cell shedding or other reasons, it will not affect the test results.

Using vortex shaking can better resuspend the cells. Vigorous vortex shaking will affect the cells that have bound antibodies and reduce the detection value. But in the actual test, it was found that the short-term vortex oscillation would not have a significant impact on the test value. The reason may be that the Gp64 antibody has a higher affinity for the antigen. The FCM method has good reproducibility for detecting virus titer. In addition to FCM’s ability to accurately recognize the signal of a single cell, the specific binding of Gp64 antibody to Gp64 also plays a decisive role. Inactivation of FBS in diluent can reduce non-specific adsorption. Due to the good specificity of binding of GP64 antibody to antigen, the content of inactivated FBS had no significant effect on the detected value in the experiment.

The cell growth status had a significant effect on the titer value. The operational process of cell passage and the cell cycle have great influence on the cell growth state, which obviously affects the titer detection value. In the operational process of multiple successive passage, the cell state is prone to decline, which is not suitable for titer detection. The poor cell growth status can be adjusted by more rigorous cell passage operation, which is suitable for titer detection again.

The grow cycle of Sf9 cell have four stages: adaptive phase, logarithmic growth phase (early, middle, and late), plateau phase, and recession phase.The density after passage has an effect on the time of the cells to enter the logarithm phase. Generally speaking, when the density is 1.0∼1.2×10^6^cells/ml after passage, the cells can enter the logarithmic growth phase within 24 hours. When the passage density is 0.8∼1.0×10^6^cells/ml, the adaptive phase is longer, and it will enter the logarithmic growth phase after more than 24 hours. The cell state in the early phase of the logarithm is the best, which is suitable for titer detection.The detection values were lower when using mid logarithmic and late logarithmic cells.The cells of recession phase are not suitable for titer detection.

The assay is based on detection of the baculovirus gp64 fusion protein which is expressed on the surface of infected insect cells after infection. Ensure that the incubation time is between 13-22 hours, which is very important to obtain accurate titer detection values. If the incubation time is less than 13 hours, the GP64 protein cannot be adequately expressed and distributed on the surface of infected cells, resulting in low detection value. If the incubation time is more than 22 hours, secondary infection will occur (the progeny virus produced first, which then infects healthy cells). In addition, the doubling time of insect cells It is 16 to 24 hours. If the culture time is much longer than the doubling time, the uninfected cells will expand, resulting in a noticeable change in the total number of living cells in the culture well, which will also affect the detection value.

